# Perceptual Expertise and Attention: An Exploration using Deep Neural Networks

**DOI:** 10.1101/2024.10.15.617743

**Authors:** Soukhin Das, G.R. Mangun, Mingzhou Ding

**Affiliations:** Center for Mind and Brain, University of California, Davis; Department of Psychology, University of California, Davis; Pruitt Family Department of Biomedical Engineering, University of Florida; Department of Neurology, University of California, Davis

## Abstract

Perceptual expertise and attention are two important factors that enable superior object recognition and task performance. While expertise enhances knowledge and provides a holistic understanding of the environment, attention allows us to selectively focus on task-related information and suppress distraction. It has been suggested that attention operates differently in experts and in novices, but much remains unknown. This study investigates the relationship between perceptual expertise and attention using convolutional neural networks (CNNs), which are shown to be good models of primate visual pathways. Two CNN models were trained to become experts in either face or scene recognition, and the effect of attention on performance was evaluated in tasks involving complex stimuli, such as superimposed images containing superimposed faces and scenes. The goal was to explore how feature-based attention (FBA) influences recognition within and outside the domain of expertise of the models. We found that each model performed better in its area of expertise—and that FBA further enhanced task performance, but only within the domain of expertise, increasing performance by up to 35% in scene recognition, and 15% in face recognition. However, attention had reduced or negative effects when applied outside the models’ expertise domain. Neural unit-level analysis revealed that expertise led to stronger tuning towards category-specific features and sharper tuning curves, as reflected in greater representational dissimilarity between targets and distractors, which, in line with the biased competition model of attention, leads to enhanced performance by reducing competition. These findings highlight the critical role of neural tuning at single as well as network level neural in distinguishing the effects of attention in experts and in novices and demonstrate that CNNs can be used fruitfully as computational models for addressing neuroscience questions not practical with the empirical methods.

## Introduction

Convolutional neural networks (CNNs) are a class of deep neural networks that draw strong structural parallels with the primate visual pathway (1–5). CNNs’ functional relevance for neuroscience has also be demonstrated in recent studies that compared single neuron activity in monkeys and fMRI activities in humans with activity in CNNs, showing that there is close correspondence between layers of CNNs and the areas within the visual hierarchy (6, 7) (8, 9). Increasingly, CNNs are being used as models of primate visual processing, making possible explorations that are not practical in biological systems (e.g., lesion), generating results that inspire new questions and new empirical experimentation (10, 11) (12, 13) (14–17). The introduction of feature-based attention (FBA) in CNNs has further deepened the integration between AI-inspired neural models such as CNNs and cognitive neuroscience (1). It has been shown that object recognition in challenging settings (e.g., images where scenes and faces are superimposed) is enhanced with feature-based attention (FBA). Attention can operate both at the level of elementary features and at the object level, which are essentially higher-order collections of features. Attention may select objects based on the presence of relevant features, implying a dual role for attention mechanisms. For example, when attention is directed toward specific features, it enhances recognition of objects that are composed of these features. The goal of this study is to leverage these developments to examine the relation between attention and perceptual expertise computationally.

Perceptual expertise refers to the enhanced ability to recognize and categorize objects in a specific category and can be acquired through extensive experience and practice. It changes how objects in the category are perceived (18, 19) and represented in the visual cortex (20–22). For example, expertise in face recognition is associated with enhanced activity in the face selective area known as the fusiform face area (FFA) (23–25). Similarly, expertise in other object categories, such as birds or cars, has been shown to increase neural activity in different category-specific regions of the visual cortex, demonstrating the broad impact of experience and practice on neural processing (26, 27). More relevant for this study, expertise also enables experts to more easily attend to the salient features of objects falling within their area of expertise (28), suggesting a possible relation between expertise and selective attention. For example, during car viewing it has been observed that manipulating attention to the identity versus the location of cars had a more pronounced impact on car novices compared to experts (27). Such effects transcend category domains and have been found in such diverse domains as planes, animals, chessboards, and radiography (26, 29–32). In all instances, experts demonstrate automatic holistic processing during object recognition, outperforming novices who are often influenced by task constraints, context, and various other factors (18, 30, 33, 34). We hypothesize that enhanced effectiveness of selective attention in the domain of expertise is a key factor underlying the superior performance of experts in their domains of expertise.

Depending on the datasets and the training objectives, CNNs can become experts in recognizing different categories of images. A recent study showed that a CNN trained on the ImageNet dataset to recognize objects became proficient at object recognition but less so on face recognition, whereas a CNN trained on the VGGFace dataset to recognize faces become proficient at face recognition, but less so on object recognition (35). In another study (9), a CNN trained to recognize both scenes and objects evolved category-selective topographical units, providing a computational account for the altered neural activity in category-selective brain regions observed in experts relative to novices. What has not been addressed in these previous studies is the relation between perceptual expertise and selective attention. We address this question by considering two classes of CNNs: one that is trained to recognize scenes (Scene-expert) and another that is trained to recognize faces (Face-expert). In our study, attention was applied at both the feature and object level by biasing neural units that preferred certain categories (faces or scenes). This approach allowed us to explore how attention modulates recognition within and outside the domain of perceptual expertise. The aim was to assess the effectiveness of attention-to-objects and attention-to-faces in each of the two CNN models. We hypothesized that (1) the scene-trained CNN would be inferior at face recognition relative to scene recognition, whereas the face-trained CNN would be inferior at scene recognition relative to face recognition, and (2) that in challenging perceptual situations (e.g., superimposed stimuli), the scene-trained CNN would benefit more from feature-based attention (FBA) for scene recognition, whereas face-trained CNNs would benefit more from FBA for face recognition. The geometry, representation, and activity of population-level neural activity were examined to explore the underlying neural mechanisms.

## Results

### Overview

We used VGG16, a 16-layer deep convolutional neural network (DNN) (36), as a model of the ventral visual stream; see Figure 1A. In this model, neurons in each layer are connected to neurons in the next layer in a convolutional manner, mimicking the receptive-field based feedforward retinotopic processing of visual information in the primate ventral visual system. Two VGG16 models were independently trained to acquire expertise in either object recognition or face recognition. Specifically, the scene network was trained on the ImageNet database (37) consisting of 3.2M images where the training objective was to classify the images into 1000 object categories as accurately as possible. On the other hand, the face network was trained on the VGGFace dataset (38) consisting of 2.6M images of human faces where the training objective was to classify the images into 2622 distinct faces as accurately as possible (Figure 1A). We will refer to the network trained on ImageNet as ‘Scene-expert’ and the network pretrained on VGGFace as ‘Face-expert’. The main purpose of this study was to examine the effectiveness of feature-based attention (FBA) to faces and to scenes in the two networks when they are engaged in performing challenging scene recognition and face recognition tasks.

**Figure 1.**
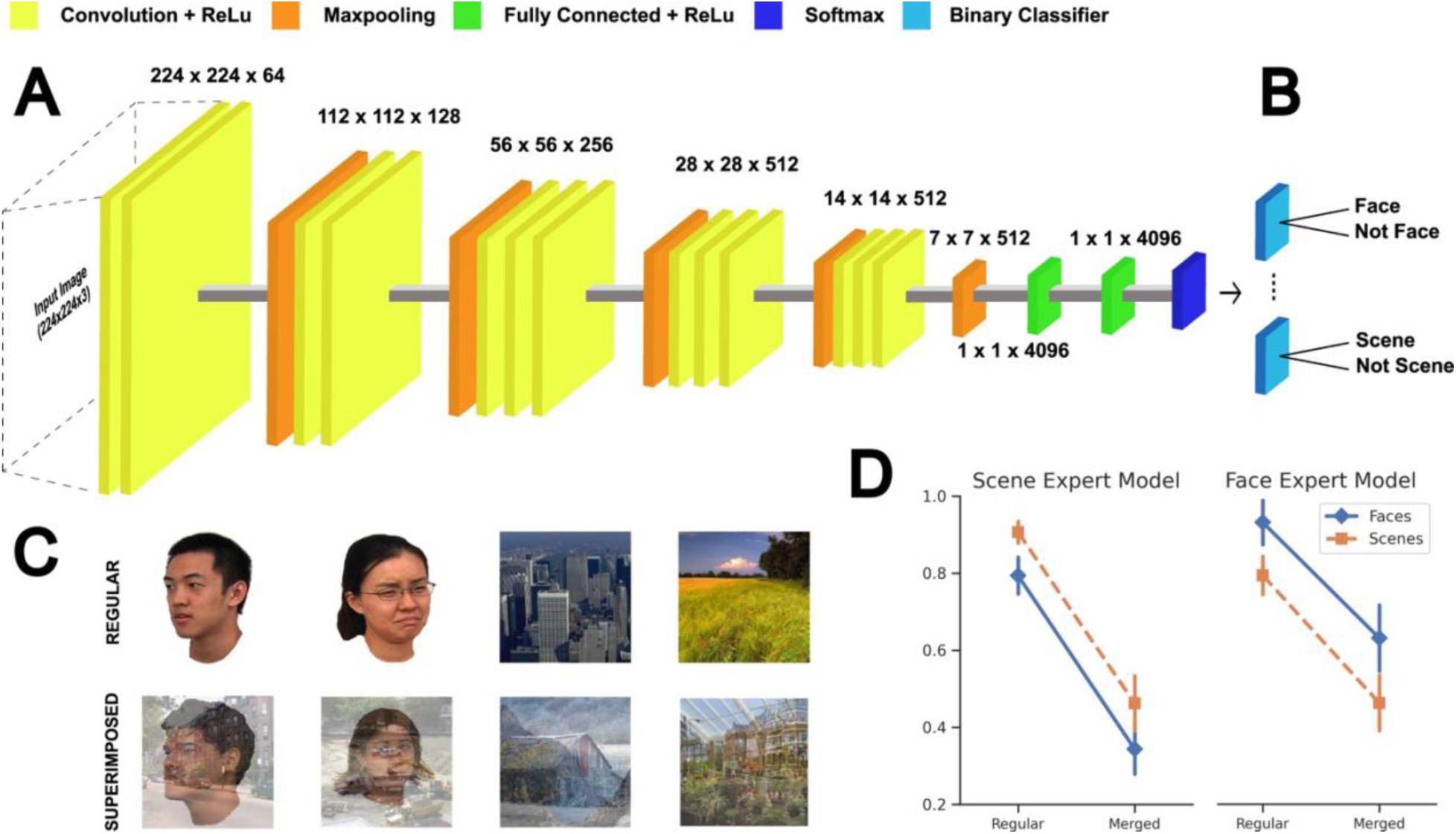
Model and study design. (A) The VGG16 convolutional neural network model (feature units and dimensions are labelled across the respective layers). One model is pretrained on ImageNet database (Scene-expert) and the other on VGGFace database (Face-expert). (B) The final layer of each network was replaced with a series of binary classifiers (logistic regression, one for each category) which were trained based on the datasets used in this study. (C) Regular and superimposed images from each category, sized at 224x224 pixels (input dimensions of VGG16). Superimposed images (bottom) were composed by transparently superimposing two images from either the same or the different categories. The regular images are used for training the binary classifiers which were then tested on both regular and superimposed images to identify the presence or absence of a certain category. (D) 5-fold cross-validation performance of the two models, image-category wise, for the Face-expert (right) and Scene-expert (left) models. Images were used from publicly available datasets (39–41).

### Experimental Paradigms and Model Performance

The recognition task consisted of identifying whether a particular object feature was present in an input image. For example, during the face recognition task, the models were asked to determine the presence/absence of a face in the input. For the model to perform these binary classification tasks, the final classification layer of the VGG16 network was replaced with a series of binary classifiers, each specific for recognizing the presence or absence of a particular object category (Figure 1B). There were two types of input: regular images and superimposed images. Regular images consisted of images belonging to one of the two categories without any distractors: faces (male and female) and scenes (manmade and natural) (Figure 1C). After training, task performance was obtained for the testing data (for more details on the training and testing dataset, see Methods). See Figure 1D. As expected, the Scene-expert model had a higher scene recognition accuracy (97.6%) for scene images compared to the Face-expert model (79.6%; p < 1e-5, paired t-test across classification folds), while the Face-expert model performed better (94%) in recognizing faces in regular images than the Scene-expert model (80%; p < 1e-4, one-sided paired t-test across classification folds). These performance metrics were summarized in Table 1. Based on these results, the two models can be said to have developed category-specific expertise, with the Scene-Expert model excelling at recognizing scenes and the Face-expert model excelling at recognizing faces.

**Table 1.** Performance of the two models for different types of images, regular and superimposed, across categories.

|  | Scene-Expert Model |  | Face-Expert Model |  |
| --- | --- | --- | --- | --- |
| Image type | Regular | Superimposed | Regular | Superimposed |
| Faces | 80% | 36.2% | 94% | 64.4% |
| Scenes | 97.6% | 48.4% | 79.6% | 48% |

For the more challenging tasks, superimposed images were utilized as stimuli; see Figure 1C. There were three types of superimposed images: (1) face over face, (2) scene over scene, and (3) face over scene (which is equivalent to scene over face). For detecting the presence or absence of a face, (1) and (3) were true positives whereas (2) were true negatives. For detecting the presence or absence of a scene, (2) and (3) were true positives whereas (1) were true negatives. The test images were balanced with 50% true positives and 50% true negatives; a total of 40 positive images and 40 negative images were used for testing. As expected, on superimposed images, the performance was significantly decreased for both models. The Scene-expert model exhibited a comparable scene recognition accuracy (48.4%) to the Face-expert model (48%; p > 0.05, one-sided paired t-test across classification folds). However, the Face-expert model performed significantly better (64.4%) in recognizing faces in superimposed images than the Scene-expert model (36.2%; p < 1e-3, one-sided paired t-test across classification folds). A summary of the performance metrics can be found in Table 1. It can be noted that both models experienced a large decrement in performance when recognizing objects in superimposed images. Thus, the presence of distractors made the task very challenging and reduced performance of each expert model, thereby presenting a fruitful opportunity to test where attentional mechanisms can be brought to bear to overcome these challenges.

### Attention Modulation of Model Performance

Feature-based attention (FBA) was implemented according to the Feature Similarity Gain Model (FSGM) (42).This model posits that neural activity is modulated in proportion to how strongly a neuron prefers an attended feature (43, 44). When a stimulus falls within the receptive field of a neuron, directing attention to that stimulus results in an increase in neural response, and this increase occurs in a proportional manner, encompassing both preferred and nonpreferred stimuli. From the foregoing, to apply FBA according to FSGM, each neuron’s selectivity for different experimental stimuli needs to be determined. This was done by calculating the responses of each feature map to the two types of stimuli: faces and scenes (see Methods). Some examples of the resulting tuning curves are shown in Figure 2. To implement FBA, the activity of a certain neuron was modulated by scaling the slope of its activation function (ReLu) in the network based on its tuning curve (i.e., how strongly a certain neural unit prefers a certain image category). Units that are selective to the attended feature (target category) had their output tuned up, while neurons that are not selective had their output tuned down (distractor category); see Figure 3. We tested the effect of FBA on one layer of each model at a time, while the performance across categories and layers that received attention modulation was recorded individually. Figure 4 and Table 2 shows the change in performances across categories for both model variants when tested with superimposed images. Overall, FBA yielded a favorable influence on model performance. Importantly, an interplay of attention’s effect across categories and models was evident upon comparative analysis. Specifically, for the Scene-expert model, attention yielded a more pronounced enhancement in performance for scenes (improvement ranging from 15 - 35% depending on the layer of the model, Figure 4A) as opposed to faces (improvement ranging from 3-15% depending on the layer of the model). In contrast, the Face-expert model exhibited significant recognition improvement on face categories (15 - 20% depending on the layer, Figure 4B) but inconsistent even negative improvement on scene categories (-12 to 10% depending on the layer). We performed a two-way analysis of variance (ANOVA) including the factors model (Scene-expert and Face-expert), layer (1 to 13) and image category (faces and scenes). The results revealed a significant main effect of model, F(1, 232) = 187.69, p < .001, a significant main effect of layer, F(12, 232) = 2.54, p = .003, and a marginally significant main effect of image, F(1, 232) = 3.29, p = .071. Furthermore, the interaction between layer and model, F(12, 232) = 6.26, p < .001, and the interaction between image and model, F(1, 232) = 636.57, p < .001, were also significant. These findings suggest that the model with perceptual expertise in each image category benefited more from FBA applied to recognize objects from that category.

**Figure 2.**
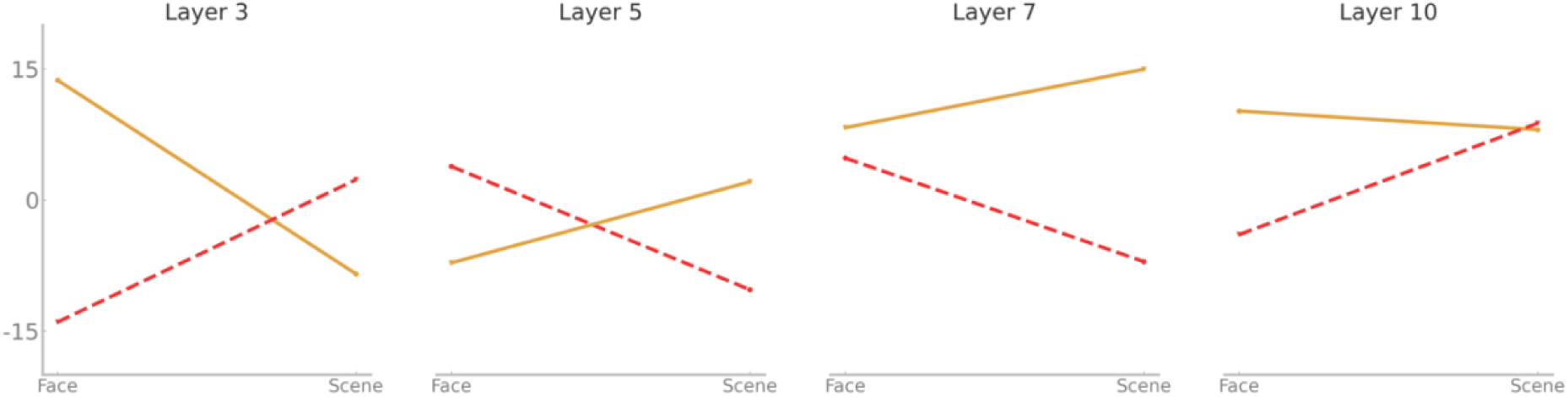
Example tuning curves of units (2 randomly chosen units are shown here) in each layer. From these tuning curves the preference of a neuron towards face or scene is determined.

**Figure 3.**
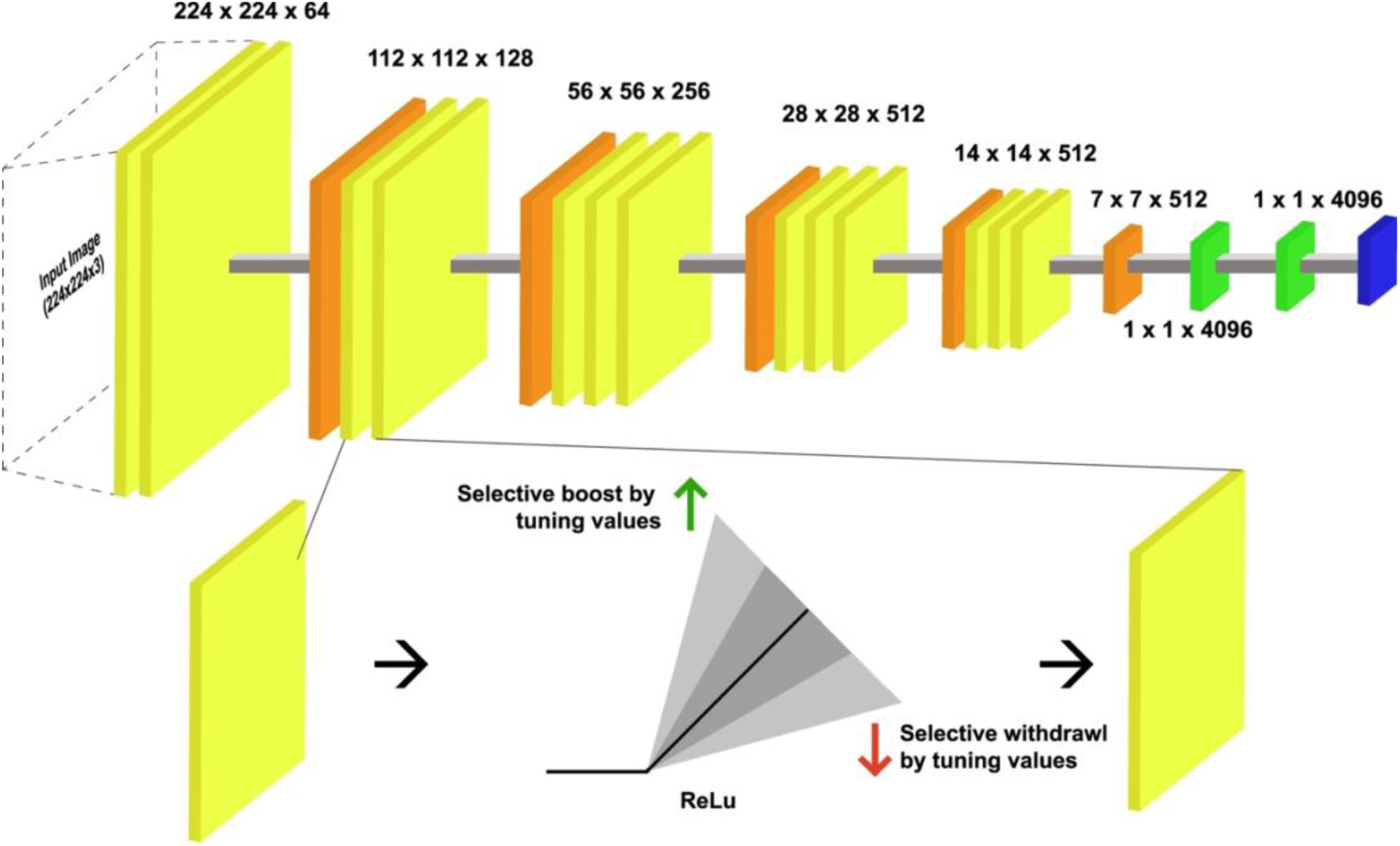
Schematic of the FBA implementation in the model. The slope of the Rectified Linear Unit (ReLu) activation function is modulated based on the tuning values of the neuron. If a certain unit in a layer prefers the attended object category, the slope of the ReLu function is tuned-up (green arrow) whereas if a unit does not prefer the attended category, its slope is tuned-down (red arrow). See the Methods section for more information about how FBA was applied.

**Figure 4.**
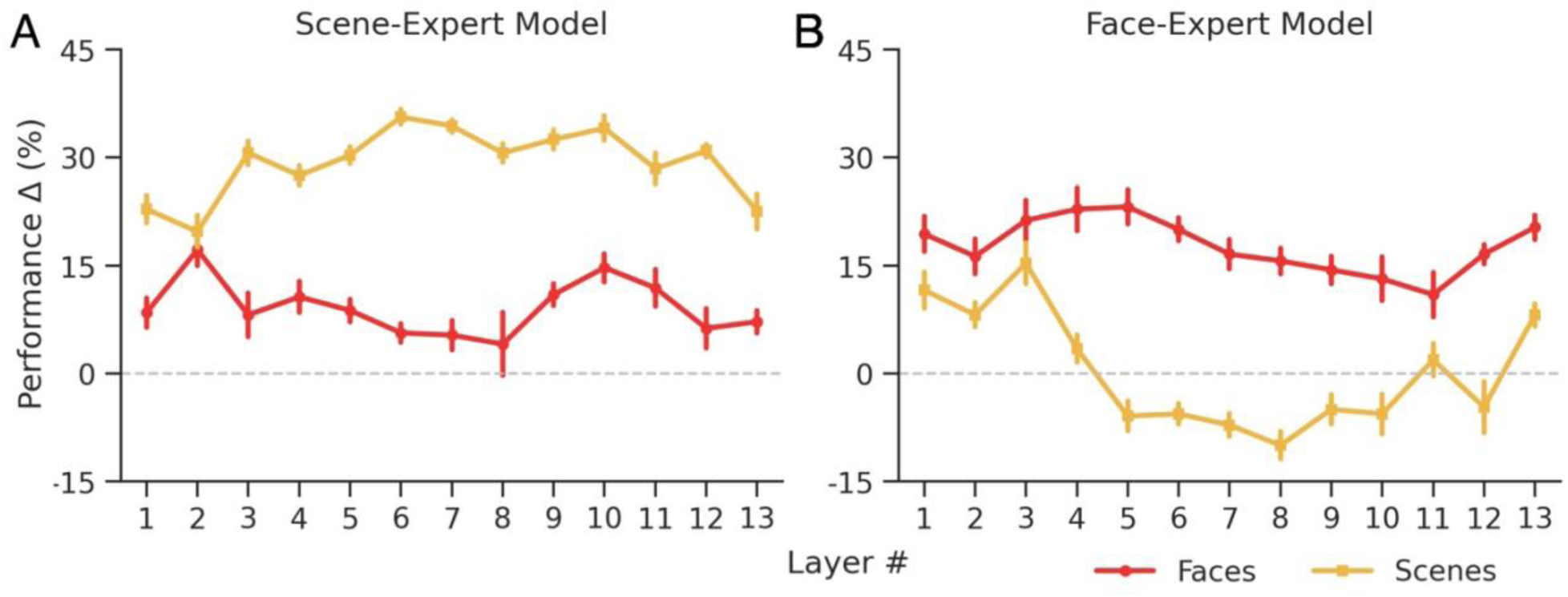
Outcomes of applying FBA to VGG16 pretrained on ImageNet (A) and VGGFace (B), across categories. Differential specificity of categories can be observed in terms of performance increases. (A) For Scene-expert model, FBA increased the performance of detecting the presence versus absence of scenes more than detecting the presence versus absence of faces. (B) For the Face-expert model, FBA was effective for enhancing the performance of detecting the presence versus absence of faces; for detecting the presence versus absence of scenes, the FBA’s effect was not very helpful, and could even be negative (i.e., decreasing the performance of the model).

**Table 2.** Summary statistics of performance improvements (i.e., Δ%) in the Face-expert and Scene-expert model across different image categories (faces and scenes) averaged across convolutional layers (1–13).

| Image Category | Model Type | M | SD |
| --- | --- | --- | --- |
| Faces | Scene-expert | 9.16 | 6.03 |
|  | Face-expert | 17.71 | 5.93 |
| Scenes | Scene-expert | 29.23 | 5.78 |
|  | Face-expert | 0.34 | 9.15 |

To ascertain the specificity of attentional modulation observed, we conducted a control experiment involving non-specific modulations of unit activity. Specifically, we applied non-specific modulations to the neurons in both models by randomly shuffling the tuning properties. For each neural unit within a given layer, we derived the tuning properties from randomly permuted sets of tuning values. We did not observe any significant increases in task performance using non-specific scaling. These results rule out the possibility of non-specific interactions influencing the attention-performance relationship. Instead, they affirm the facilitating influence of FBA on task performance, particularly highlighting its capacity to interact in an expertise-specific manner. Furthermore, by employing randomly permuted tuning values instead of equal scaling values (as used in (1)), our findings extend previous research. It is noteworthy that merely permuting the labels of tuning curves does not yield the same effect of attention, underscoring the effectives on FBA implemented via the FSGM.

### Potential Mechanisms of Enhanced Effectiveness of FBA in Expert Networks

Studies have shown that attention modulates a neuron’s response according to the neuron’s tuning properties in early and late layers of the visual system (45, 46). In these studies, the response of a neuron is substantially enhanced when an optimal stimulus is attended, whereas its response to an attended non-optimal stimulus is enhanced to a lesser extent, or even decreased (47, 48). So, we probed if there was a similar effect of attention on neurons’ tuning properties in our models, and if it was related to expertise. For this, we analyzed the tuning curve of each neuron and computed its tuning quality, which was its maximum value. Tuning quality provided a quantifiable measure of the strength with which a particular neuron exhibited preference for a specific object category. This was done separately for scene- and face-selective neurons, and then compared to each other. The distribution of tuning quality across layers was analyzed during the baseline condition (without attention), and when attention was applied to neural units that preferred the target category for each layer individually. This enabled us to investigate whether there was any expertise and/or image category preference-related differences that resulted in differential effects of attention in the previous behavioral analysis. Figures 5A and 5B show the results without attention (see Methods for a description of the statistical tests). During the baseline condition, neurons in the Scene-expert model demonstrated a significantly stronger preference for scenes, as evidenced by the higher tuning quality for scenes compared to faces (p < 0.001 for layers 3 to 12, one-sided Welch’s t-tests across layers, FDR corrected for multiple comparisons). Conversely, the Face-expert model exhibited significantly greater tuning quality for faces compared to scenes, suggesting a stronger preference for faces (p < 0.001 for all layers except layer 7, one-sided Welch’s t-tests across layers, FDR corrected for multiple comparisons).

**Figure 5.**
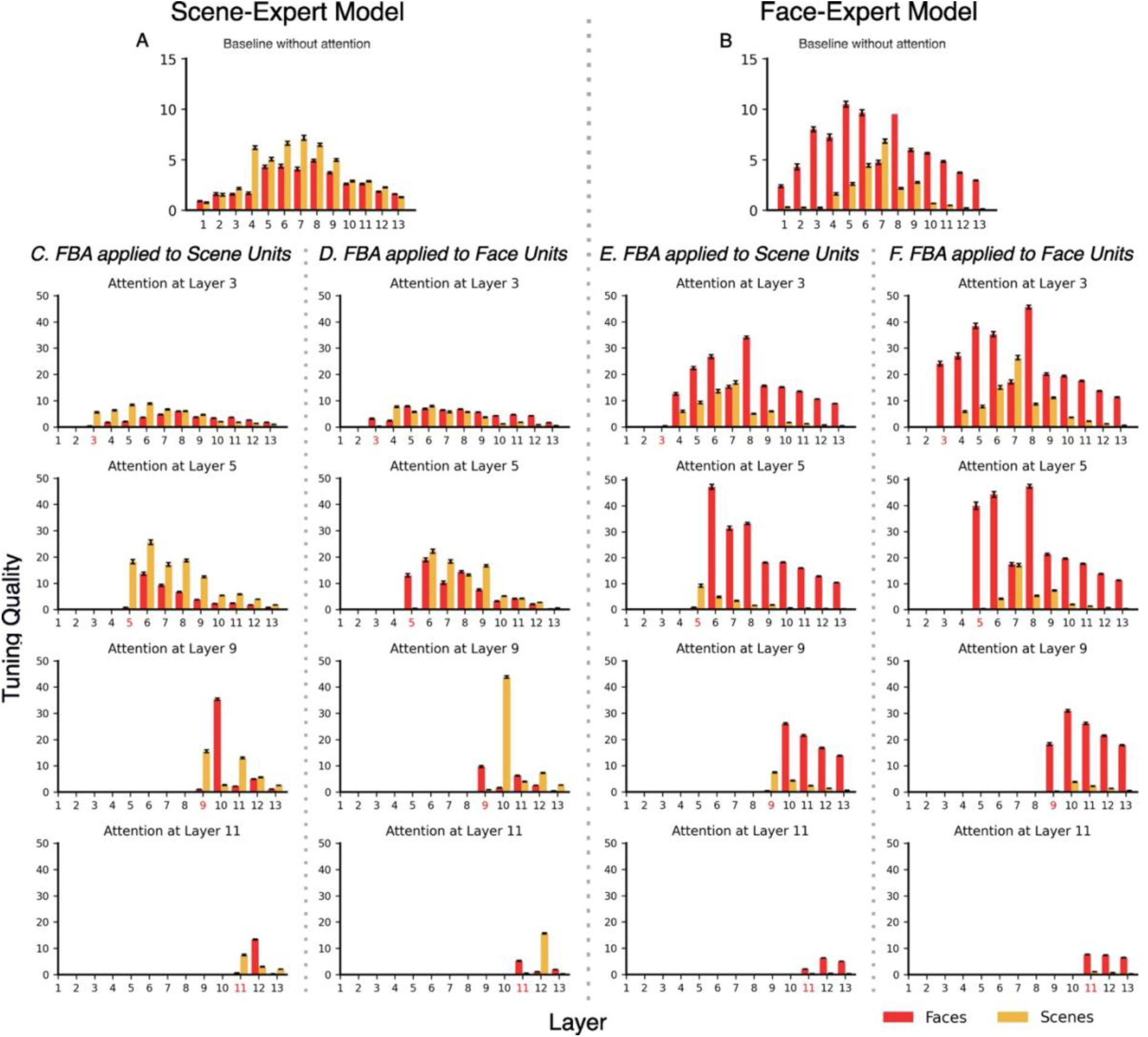
Tuning quality across layers in the Scene-expert (A) and Face-expert (B) network divided into face (red) and scene (yellow) selective neurons during baseline when attention was not applied. Bars indicate the tuning quality distribution of neurons across layers 1 through 13. Tuning quality of neurons that prefer scenes is higher than that that prefer faces in the Scene-expert Model (A) and vice-versa in the Face-expert Model (B). (C-F) Tuning quality distribution divided based on FBA applied to scene and face selective neural units when attention is applied at different layers (row-wise layers 3,5,9 & 11 shown) in the Scene-expert Model (C-D), and the Face-expert Model (E-F).

Given that feature based attention is multiplicative, for the same strength of attention modulation (i.e., the same 𝛽 value), the face-selective neurons in the Face-expert will have a stronger increase in activity than the scene-selective neurons, resulting in the higher performance improvement in detecting the presence versus absence of faces in the input. The same principle could explain the reason why in the Scene-expert case, for the same strength of attention modulation, there is a higher performance improvement for detecting the presence versus absence of scenes in the input. We examine these ideas next.

In Figures 5C - F, we show the results when attention was applied to investigate whether attention modulated the tuning quality across different image types. We analyzed the effect separately for scenes and faces in the two models while attention was applied at different layers, showing representative results from layers 3, 5, 9, and 11 in the figure. Applying attention to different layers resulted in stronger tuning quality of neural units compared to baseline, in manner that is in accordance with the principles of the FSGM model. However, the degree of modulation varied across models and image categories. In the Scene-expert model, as illustrated in Figure 5C, applying attention separately to scene-selective units enhanced their tuning quality (yellow bars), surpassing that of face-selective units (red bars). This effect began at the layer where attention was applied and persisted through higher layers, maintaining the higher tuning quality of scene-selective units throughout the model. In contrast, when attention was directed to face-selective units (Figure 5D), their tuning quality (red bars) did not exceed that of scene-selective units (yellow bars). Although face-selective units experienced improvement in tuning quality at the layer where attention was applied (in line with the FSGM principle), this enhancement did not sustain or carry over to higher layers, unlike the consistent propagation observed for scene-selective units. Similarly, in the Face-expert model (Figure 5E), attention selectively improved the tuning quality of face-selective units, but this effect did not significantly influence other types of units (Figure 5F). Thus, in each expert model, the tuning quality of neurons was enhanced by attention only when their preference aligned with the category of expertise specific to the model. Conversely, attention did not improve the tuning quality of neurons whose category preference differed from the model’s expertise category.

### Representational Similarity Reveals Feature Separation in Models

In the previous section, we examined the interplay between effects of attention and expertise-specific task performance improvement by analyzing the quality of neuron-level tuning in our models. However, apart from competition and bias that has been observed at single-unit level, the expertise-attention interaction may also arise due to differences in network-wide neural representations. Prior research has demonstrated that top-down bias and task performance is context dependent, with greater competition occurring between stimuli that are more similar or closer to each other in terms of neural representation (47, 49). Therefore, we investigated the impact of dissimilarity in neural representations of different categories on the underlying effects of attentional bias.

First, we applied representational similarity analysis (RSA) to assess the degree of dissimilarity and competition between different image types (faces and scenes) across layers of each model, taking into account the model’s domain of expertise. Next, we conducted the same analysis on images when FBA was applied to neurons in different layers, selectively targeting either face- or scene-preferring neurons. This approach enabled us to examine how attention modulates the multivariate neural representation of each image type within its corresponding expert model. Finally, we compared the effect of attention on RSA dissimilarity and competition between face and scene representations within each model, highlighting how attentional mechanisms interact with expertise to shape multivariate neural representations.

The representational dissimilarity matrices (RDMs) for face-expert and scene-expert models across different layers are shown in Figure 6A (top-left panel). The patterns of dissimilarity change across layers, indicating that the models process and differentiate each image category from the other image category at each processing stage (layer) of the network. A theoretical RDM, as illustrated in Figure 6B, represents an idealized categorical structure, where two categories of images (faces and scenes) are perfectly separated. It serves as a reference for understanding the relation between face and scene representations in the two models. We assessed the relation between RDMs obtained from each layer in the two models (baseline without attention) and the theoretical RDM using rank-ordered Spearman correlations. This yielded 26 (13 layers x 2 models) RSA correlation measures, each depicting the degree of dissimilarity or separation between face and scene features encoded in the model layers (Figure 6C shows mean RSA correlation values with ±1 SEM, bootstrapped across 100 iterations, baseline without attention). Throughout the layers, we observed modest dissimilarity measures (0.1 ± 0.03) except for early layers in the Face-expert model, which exhibit higher dissimilarity between the features. These values are reasonable and within the bounds of what has been reported in similar studies (50, 51).

**Figure 6.**
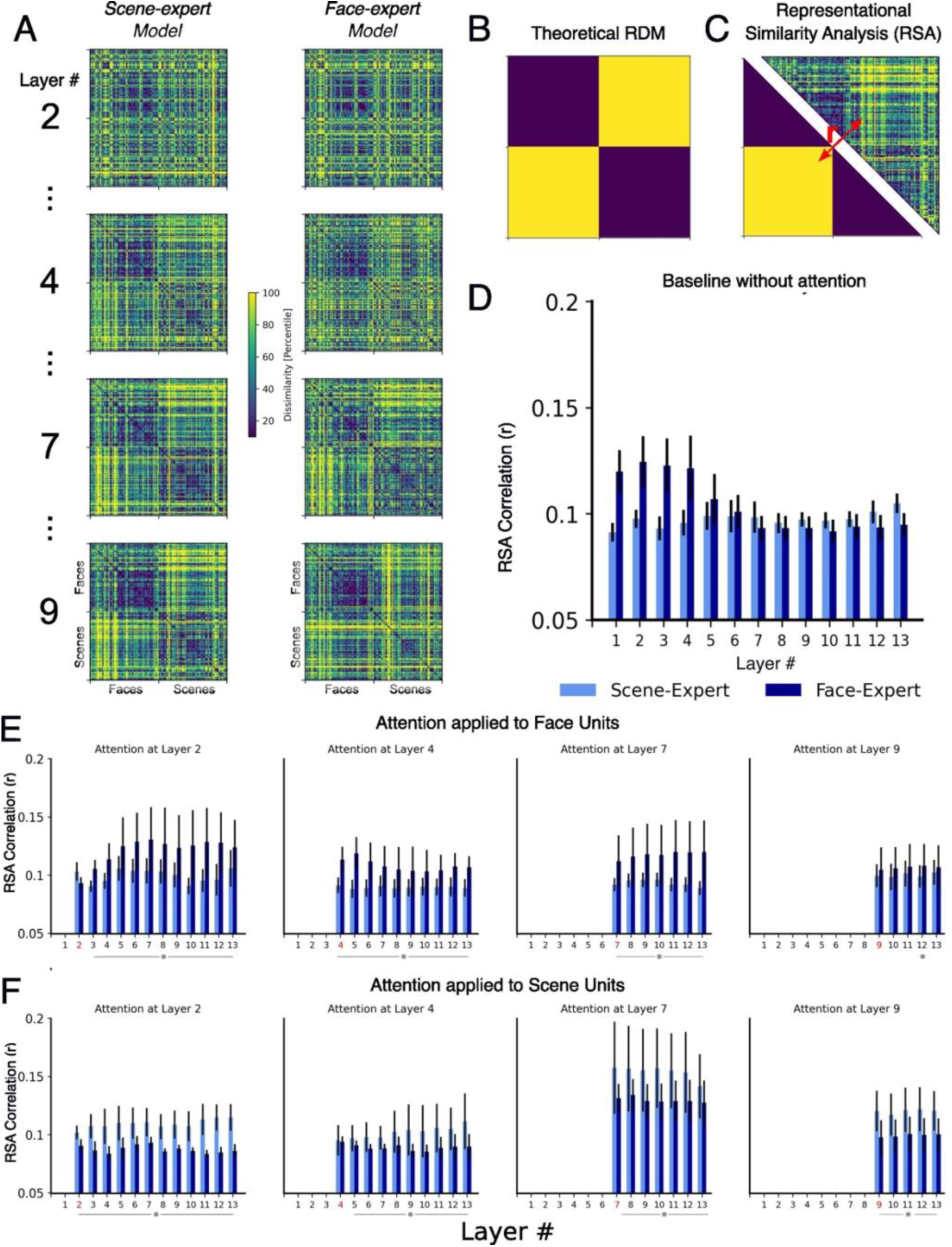
Representational Similarity Analysis (RSA) across different categories of images and models. (A) Representational dissimilarity matrices (RDMs). For each layer within a model, separate RDMs were constructed for all scene and face images by one minus the Pearson r correlation between each pair of image-evoked multivariate neural activations. Here, RDMs are shown for layer 2, 5, 9 and 12 for each model. (B) Theoretical RDM representing the ideal degree of separation between Scene (Manmade and Natural) and Face (Male and Female) images. (C) RSA analysis, performed by calculating the rank-ordered Spearman correlation between the off-diagonal triangular values of the theoretical RDM and layer RDMs. (D) RSA analysis for each model, layer-wise during the baseline condition when attention was not applied to any neural units in Scene-expert (light blue) and Face-expert (deep blue) models. (E, F) Representational Similarity (in Spearman rho correlation) when attention was applied to Face-selective units (E) and Scene-selective units (F) at layers 2,4,7 & 9 (left to right, highlighted with red layer labels on the x-axis). Error bars indicate ± 1 SEM obtained from bootstrapping technique using 100 samples. * = p < 0.05, one-tailed paired t tests and FDR corrected for multiple comparison across layers.

Next, we computed the relation between theoretical RDM and the data based RDMs from each layer (in a model) when attention was separately applied to neural units that either prefer faces (Figure 6E) or scenes(Figure 6F). The mean RSA correlation values, along with ±1 SEM, were bootstrapped across 100 iterations and shown for layers 2, 4, 7, and 9 as representative examples. In both models, we observed consistently positive mean RSA correlations across all layers, with representational similarity patterns varying based on model expertise and the type of neurons to which attention was applied. When attention was applied to face-selective neurons, the similarity values were higher in the Face-expert model compared to the Scene-expert model. This indicates the effectiveness of attentional mechanisms in inducing greater separability between face representations and scene representations in the Face-expert model (p < 0.05, one-tailed paired t-tests, FDR corrected for multiple comparisons across layers). Similarly, when attention was applied to scene-selective neurons, RSA correlation values were greater in the Scene-expert model compared to the Face-expert model (p < 0.05, one-tailed paired t-tests, FDR corrected for multiple comparisons across layers). Therefore, the Scene-expert model was more efficient in distinguishing scenes from faces when attention was applied to scene-selective neurons only. Similarly, the Face-expert model was able to identify faces from scenes efficiently when attention was applied to face-selective neurons only.

The expertise in each model allows for finer discrimination in the multivariate neural representations between object features belonging to their category of expertise. Since the effectiveness of attention depends on the neural representation of stimulus features, which is in turn dependent on the expertise of the model, it is more effective when domain-specific objects are distinctly encoded in various layers. This finding supports the feature similarity gain model of attention, demonstrating that attention selectively enhances the processing of features relevant to the model’s expertise.

## Discussion

In this work, we investigated whether and how perceptual expertise interacts with the effect of attention in a convolutional neural network (CNN) model of the primate ventral visual system. Specifically, we used neuron-level tuning and multi-neuron network-wide neural representation to examine the mechanisms underlying the observed interaction between model expertise and top-down attention. Our results complement the previous neuroimaging and electrophysiological studies suggesting that top-down attention control interacts with neural pathways and brain regions associated with perceptual expertise and enhance performance in an expertise-dependent fashion (27, 29, 52–54).

### The interplay of expertise and attentional bias

We trained a VGG16 neural network model to specialize in recognizing either scenes or faces. The Scene-expert model was trained with scene images, while the Face-expert model was trained with face images. During the experiment, the images were presented in two forms: regular images, which were single images from a category (faces or scenes), and superimposed images, which were superimposed images from the same or different categories. The results established the presence of category-based expertise in the models. Specifically, the Face-expert model demonstrated superior performance in recognizing the presence or absence of faces in the single image input, while the Scene-expert model excelled at detecting the presence or absence of scenes in the single image input. When presented with superimposed images, the recognition performance declined significantly, indicating that the models were not effective at dealing with distractors in the superimposed images. Next, we applied feature-based attention to neurons based on their tuning profiles. The Scene-expert model showed significantly larger improvements in recognition performance with attention when detecting the presence vs absence of scenes as compared to detecting the presence versus absence of faces. Similarly, the Face-expert model showed significant improvement with attention when detecting the presence versus absence of faces than detecting the presence vs absence of scenes. Expertise in a specific category allowed the models to develop specialized neural representations that are optimized to interact with attention and improve task performance.

### Expert Versus Novice: Unit Level Analysis of Object Based Attentional Enhancement

To understand why attention is more effective when combined with expertise, we analyzed the tuning quality of artificial neurons, which reflects how strongly they prefer an object category. Our findings indicate that the Scene-expert model exhibits stronger tuning quality for scenes than for faces, while the Face-expert model shows stronger tuning quality for faces than for scenes. According to the Feature-Specific Gain Modulation (FSGM) model, attention modulates the neuron’s firing by a multiplicative factor applied to the tuning function (43, 44, 55). Given that the tuning function here is computed over the two categories of images, this operation will lead to the enhancement of the attended category (target) and the suppression of the ignored category (distractor). Given the baseline tuning quality findings mentioned above, the multiplicative nature of the attention amplification further ensures that the attention enhancement is stronger when it is directed to the image category in which the network is an expert. Thus, attention is more effective in experts because it operates on neurons that are already highly tuned towards the object of expertise. This selective tuning, in addition to already sharpened tuning due to expertise, likely underlie the enhanced effect of attention when the attended object matches the expertise of a model. Neurophysiological studies of experts and novices consistently report this phenomenon, highlighting how experts often exhibit increased activation levels within brain regions relevant to their task and sometimes recruit additional areas involved in domain-specific processing (23, 26, 56, 57). For instance, Gauthier et al. (1999) found that car experts showed greater activation in the fusiform face area (FFA) when recognizing cars compared to novices, indicating specialized processing. Similarly, Maguire et al. (2002) observed that London taxi drivers, who are experts in navigation, had larger hippocampal volumes and showed increased activation in this region during navigational tasks. Grabner et al. (2006) reported that individuals with high mathematical expertise exhibited greater activation in brain areas associated with arithmetic processing, and McGugin et al. (2014) found that bird and car experts showed enhanced activation in category-specific regions when recognizing birds and cars, respectively. These studies collectively suggest that expertise enhances the neural efficiency and sharpens the tuning curve in favor of features that match its domain of expertise.

### Population Level Analysis: Representational Similarity of Targets and Distractors

It has become increasingly clear that visual perception relies on multivariate representations of visual inputs. Stronger separation between the neural representations of targets and distractors is crucial for the effectiveness of attentional selection of task-related stimulus information. Previous studies have demonstrated that efficiency of these mechanisms is heavily influenced by individual differences in the representation of task-related information (58–61). Specifically, unique category representations, formed through top-down attentional templates, are pivotal in guiding the search for targets and suppressing distractors. These templates are shaped by an individual’s experience and expertise, meaning that a person’s ability to effectively utilize attentional mechanisms depends on how well their mental representations align with the task at hand.

According to the biased competition model of attention, visual stimuli compete for neural representation in the brain, and the more similar the targets and the distractors, the stronger the competition (47, 62, 63). Conversely, the more dissimilar the targets and the distractors, the weaker the competition, the better the behavioral performance. Our results are consistent with this. In our study, Representational Similarity Analysis (RSA) revealed that at the multivariate neural representational level, attentional mechanisms are effective primarily for object categories that align with the expertise of a model. When attention is directed towards neurons within a specific layer of a model, this focus sharpens population neural tuning by increasing the representational distance between targets and distractors. As the neural features of targets and distractors become more dissimilar, the competition between them diminishes, thereby enhancing the potency of attentional mechanisms.

Recent work by Doostani et al. (47) has shown that sharpened neuronal tuning amplifies the influence of top-down mechanisms, especially in difficult scenarios where targets and distractors share common features. Our findings offer direct evidence on the strength of the attentional bias towards the target such that it increases as neuronal tuning sharpens. Our findings indicate that this effect is more significant in models with expertise in specific object categories, as expertise increases the dissimilarity between targets and distractors, thereby sharpening the tuning curve. This implies that specialized knowledge enhances the ability of attentional mechanisms to distinguish between similar features.

### Relation with Prior Literature

The association between perceptual expertise and attention mechanisms has been studied in the past. It has been reported that expertise has a facilitatory effect on categorization and involves deployment of top-down mechanisms to engage object processing as: (1) Engagement of attention transcends lower-level features of the object and prioritizes them based on their content (64, 65). Bukach et al. (2006) demonstrated that experts in face recognition could categorize faces more efficiently than novices, suggesting that expertise enhances the ability to process complex visual stimuli. Similarly, van der Linden et al. (2014) found that expert radiologists could detect abnormalities in medical images more accurately, indicating that expertise involves sophisticated attentional mechanisms. (2) Expertise entails top-down activity crucial for evaluating and recognizing pertinent stimulus features (29, 66). Harel et al. (2010) showed that experts in visual search tasks could quickly identify target objects among distractors, highlighting the role of top-down processes in expertise. Reddy et al. (2007) found that expert athletes could anticipate the actions of opponents more effectively, demonstrating the importance of top-down attention in dynamic environments. (3) Studies have also shown that experts exhibit differences in attentional selection and interacting with objects that are relevant to them. Stokes (2021) reported that expert musicians could focus on relevant musical elements more efficiently than novices, indicating differences in attentional selection. Kundel et al. (2007) found that expert radiologists took less time to fixate on abnormalities in medical images, suggesting that expertise leads to faster identification of relevant features. Vogt & Magnussen (2016) observed that expert chess players had shorter saccades and fewer fixations when analyzing chess positions, indicating more efficient visual processing. This suggests that experts possess a rapid sensitivity to holistic features of the stimulus array, indicating that their attentional processes are more efficient and diagnostically relevant to the task. Our findings extend upon prior work, by delving into the intricate interplay between biologically inspired attention mechanisms and the intricate architecture of neural networks, and their predisposition to knowledge. Also, our work helps by aligning network behaviors with established neural processes, making it easier to decipher the rationale behind network decisions, fostering robustness, transparency, and development of more interpretable DNNs.

## Summary

This study was aimed at understanding the association between attention mechanisms and perceptual expertise. We used deep neural networks as models of the ventral visual pathway of the brain for this purpose. Our findings indicate that when a deep neural network is subjected to complex tasks, its performance can be enhanced by introducing attentional mechanisms, and the effectiveness of this attention enhancement is closely tied to the perceptual expertise developed in the model, i.e., attention is more effective in a model when the task is within the model’s domain of expertise. Mechanisms at the individual neuronal response level and at the neural population level were investigated. Methodologically, investigating neural mechanisms of perception in deep learning models represents a convergence of AI and neuroscience, offering (1) a computational platform in which manipulations not practical in empirical experiments can be carried out and (2) the potential to build more efficient, adaptive, and human-like artificial intelligence systems. As our understanding of both artificial and biological neural networks advances, we can expect more refined and effective models to emerge that can better bridge the gap between artificial and natural intelligence.

## Methods

### The Model

VGG16 is used as a model of the ventral visual stream (36). VGG16, as shown in Figure 1A, is a feed forward convolutional neural network with 13 convolutional layers followed by 3 fully connected layers. In this work, two pretrained VGG 16 models were considered: one pretrained on the ImageNet dataset and thus an expert in object recognition and the other pretrained on the VGGFace dataset and was thus an expert in face recognition (37, 38). For the model trained on the ImageNet dataset, the last layer of the network outputted the labels of 1000 object categories (92.7% accuracy), whereas for the model trained on the VGGFace dataset, the last layer outputted the labels of 2622 individual faces (98.95% accuracy). In this work, the network weights connecting the first 15 layers of the VGG 16 models were taken from (36) and from (38), respectively, and kept unchanged (frozen). The last layer was replaced by a layer with output suitable for our paradigms (see below). The weights connecting the paradigm-specific last layer and the layer preceding it were trained using the datasets described below according to the goals of the paradigm.

### Image Category Detection Task

The models were presented with images containing objects from two categories: faces (male faces & female faces) and scenes (natural scenes & manmade scenes). The model’s task was to identify in the image the presence or absence of objects from one out of the two categories (e.g., is there a face in the image?). To accomplish this binary classification, we replaced the final SoftMax layer of the original pretrained VGG16 models with a layer containing a series of binary classifiers, with each classifier consisting of two units capable of signaling the “presence” or “absence” of the to-be-detected category (Figure 1B). Specifically, the Face-expert has two associated binary classifiers with one detecting the presence and absence of a face in the input and one detecting the presence and absence of a scene in the input, and the Scene-expert also has two associated binary classifiers that perform the same tasks.

To train the binary classifiers, we used a separate set of training and testing data, sourced from different image repositories (39–41), which did not overlap with ImageNet or VGGFace datasets. The dataset consisted of 200 faces and 200 scenes (224x224 pixel RGB images); see Figure 1C for examples. We used 160 faces and 160 scenes for training and the remaining 40 faces and 40 scenes for testing. During the training of the binary classifiers, depending on the category to be detected, there were always 160 true positives from the category along with 160 true negatives from the other category. The classification performances reported here were achieved by implementing logistic regression in the binary classifiers.

Once the binary classifiers were trained, they were tested separately on regular as well as superimposed images. When the input to the network model was a regular image, the network’s task was the same as the task it was trained on, namely, to detect the presence or absence of a particular category in the input image. To challenge the model, besides using regular images, we introduced superimposed images to make the task more difficult. The superimposed images were created by transparently superimposing two images either from the same category or from different categories (by taking the mathematical average of the corresponding pixels of the two images). There were three types of superimposed images: (1) face over face, (2) scene over scene, and (3) face over scene (which is equivalent to scene over face). For detecting the presence or absence of a face, (1) and (3) were true positives whereas (2) were true negatives. For detecting the presence or absence of a scene, (2) and (3) were true positives whereas (1) were true negatives. The test images were balanced with 50% true positives and 50% true negatives; a total of 40 positive images and 40 negative images were used for testing.

It is expected that the detection accuracy for superimposed images would decline significantly relative to regular images. This scenario then provided us the opportunity to apply feature-based attention to enhance the detection performance and test whether such enhancement depends on the expertise of the network. We expected that for the Face-expert, applying attention to face would enhance face detection performance more than applying attention to scene would scene detection performance, and that for the Scene-expert, the opposite is true.

### Attention Modulation of Neuronal Responses

The main goal of this study was to examine how attention interacts with perceptual expertise in a deep neural network model of the ventral visual stream. Specifically, we implemented the feature similarity gain modulation (FSGM) model of feature-based attention, following the procedures of (1). To apply this attention mechanism, we calculated the extent to which each neuron in a model (Scene-expert or Face-expert) preferred a certain object category, i.e., their tuning values. Here the term neuron was used to refer to a filter or a feature map in the model. Since feature-based attention is a spatially global phenomenon (1, 67), the responses from all the neural units within a filter were averaged to become the response of the filter or neuron. We implemented attention by modulating the slope of the ReLu function of the units within a filter according to the filter’s tuning function.

### Calculation of Tuning Values

To determine the tuning function of each neuron, we presented the model with the regular images from the two categories (the same set of images of 160 faces and 160 scenes) used for training the binary classifiers) and measured the relative activity levels of the units within the filter. As indicated above, considering that feature-based attention is nonspatial (1, 67), we treated the activity levels of all units within a filter identically and calculate its tuning by z-scoring their activity across categories using the following equations:

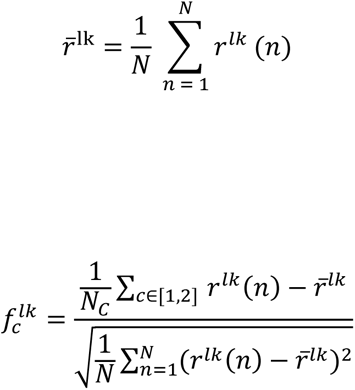

Specifically, for the *k*^th^ neuron in the *l*^th^ layer, *r^lk^(n)* is defined as the mean activity of all units in the filter in response to *n^th^* image. Finally, by taking the mean across all these values for the training set images (N_c_ = 160 images per category, c = c^th^ category with c = 1 or 2 because there are 2 categories in total; N = 2 x 160 = 320), we get the mean activity of the neuron *r̄^lk^*. The tuning value for each neuron for a given category is the z-scored mean activity with respect to the mean activity of the unit for all images. Put it simply, tuning value for a certain category for a neural unit is the average activity of the unit in response to the images from the category subtracting the mean activity across all images and divided by the standard deviation of the activity across all images. Calculated across the two categories, we get a 2-dimensional vector of values 𝑓_*c*_^𝑙𝑘^, which is the tuning function of each neuron used to implement attention. To find the preferred category of a neural unit, we designate the category with the larger tuning value as its preferred category. Tuning quality for a certain neuron is defined as the maximum tuning value of that neural unit i.e., max (|𝑓_*c*_^𝑙𝑘^|). Tuning quality is a measure of the extent of how strong a certain neuron prefers its most preferred category.

### Implementation of Feature Based Attention

We implemented FSGM attention model in its multiplicative and bidirectional form across the layers in the two networks. To apply attention at a unit from neuron *k* in layer *l* and for category c, we modulated the slope of the corresponding rectified linear unit (ReLu) by the tuning value of that category c, weighted by a strength parameter ß (varied from 0 to 20 in increments of 0.1.

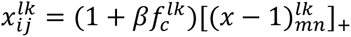

where 𝑥*_ij_*^𝑙𝑘^is the unit response at *(i, j)^th^* spatial location in the *k^th^* neuron of the *l^th^* layer, []_+_ is the ReLu function. (𝑥 − 1)*_mn_*^𝑙𝑘^ represents the activity of the *(m,n)^th^*unit from the preceding layer.

### Tuning Quality Analysis

As mentioned previously, tuning quality was defined as the magnitude of the maximum tuning value of a neuron: max (|𝑓_*c*_^𝑙𝑘^|). Consequently, it was a measure of the relative strength of the neuron’s preference towards its favored object category. It is reasonable to expect that for the Face-expert, the tuning quality for face stimuli will be higher than that for scene stimuli, whereas for the Scene-expert, the tuning quality for scene stimuli will be higher than that for face stimuli. For quantitative analysis, since there were unequal numbers of neurons selective to each category in a layer, we performed a comprehensive statistical assessment. This involved subjecting the layer-wise tuning quality distributions of the two models focusing on the categories – faces and scenes, to Shapiro-Wilk normality and Levane’s variance tests. The results of the Shapiro-Wilk test revealed a substantial deviation from the assumption of normality, consistently observed across layers (except layers 1-3 and layer 13) and model variations (p < 0.001). Furthermore, acknowledging this deviation, Levane’s test indicated the presence of uneven variances in both distributions (p < 0.001, spanning all layers in both models). Therefore, the two distributions were compared using Welch’s t-tests, FDR corrected for multiple comparisons.

### Representational Similarity Analysis (RSA)

For each model and its individual layers, we calculated the output activity of the neurons present in each layer for every image. These neural representations were then analyzed using a Representational Similarity Analysis (RSA) that consisted of two primary steps. Firstly, we generated separate Representational Dissimilarity Matrices (RDMs) for each model type (Figure 6A), obtaining one RDM per layer. These matrices were based on the distinct patterns of activation that each image elicited, categorized separately for each object category. Secondly, employing a theoretical RDM illustrated in Figure 6B, we captured the idealized maximum possible divergence between the two categories. Lastly, we computed the representational similarity by calculating the rank-ordered Spearman correlation between the theoretical RDM and the RDM for each layer of the VGG16 model.

For each model, we subjected it to all possible regular images and extracted the activation patterns across all neurons for each layer. This was done separately for each face (total=160) and scene image (total=160). The activation pattern for each image was then transformed into a one-dimensional vector. To assess dissimilarity between these vectors, we calculated one minus the Pearson’s correlation coefficient for every pair of vectors. This process resulted in the generation of 320 x 320 sized Representational Dissimilarity Matrices (RDMs) for each layer, model variant, and image category (face and scene), amounting to a total of 13 x 2 matrices. Within each RDM, the cells contained dissimilarity values (1 - *r*) representing the dissimilarity in neural representations between pairs of images. Subsequently, a theoretical RDM was constructed to match the dimensions of the VGG16 model layer’s RDMs (320 x 320). In this theoretical RDM, the cells in the top-left 160 x 160 and bottom-right 160 x 160 sections along the diagonal were assigned a value of 0, while all other cells were assigned a value of 1 (Figure 6B). This configuration indicated minimum dissimilarity (0) for images within the same category (e.g., face vs face or scene vs scene), and maximum dissimilarity (1) for images belonging to different categories (e.g., face vs. scene or scene vs. face). This method allowed for a systematic comparison of dissimilarity between different categories and facilitated the evaluation of the model’s ability to distinguish between faces and scenes at various layers.

Finally, representational similarities were computed as the rank-ordered Spearman correlation between each RDM from layers of the two models and the theoretical RDM. This process resulted in a set of similarity values (13x2), corresponding to each layer and model of expertise. To cross validate across our test set, we used a bootstrapping technique to assess the statistical significance of these values. We generated 100 test image sets with replacement, recalculating representational similarity for each sample. This process yielded an empirical distribution of these values, along with bootstrapped mean estimations and 95% confidence intervals. To identify significant differences in the mean similarity values across object categories, we employed a p-value threshold of 0.05. We rejected the null hypothesis if the confidence interval did not include 0. Furthermore, we subjected the results across layers to FDR correction to account for multiple comparisons across layers.

### RSA with attention

We repeated the same RSA analysis procedure described above when attention was applied at each layer individually to neurons that prefer faces or scenes (Figure 6E & F). Specifically, attention was modeled by amplifying the activation of specific units that exhibited a stronger response to either faces or scenes within each layer of a model (separately for the scene-expert and face-expert model). The attention-modulated activations were then processed similarly to the unmodulated activations: we calculated the output activity of each regular image, generated RDMs, and performed RSA by correlating these attention-modulated RDMs with the theoretical RDM.

## Acknowledgements

This work was supported by NIH grant MH117991, NSF grant BCS2318886, and NSF grant BCS2318984. We are grateful to Joy Geng, John Henderson, Ruogu Fang, Randall O’Reilly, Lee Miller, and the members of our labs for their helpful comments and advice.

## References

1. Lindsay GW, Miller KD. How biological attention mechanisms improve task performance in a large-scale visual system model. eLife 2018. p. 1–29.

2. Xu K, Ba J, Kiros R, Cho K, Courville A, Salakhudinov R, et al. Show, Attend and Tell: Neural Image Caption Generation with Visual Attention. In: Francis B, David B, editors. Proceedings of the 32nd International Conference on Machine Learning; Proceedings of Machine Learning Research: PMLR; 2015. p. 2048–57.

3. Cao C, Liu X, Yang Y, Yu Y, Wang J, Wang Z, et al. Look and Think Twice: Capturing Top-Down Visual Attention with Feedback Convolutional Neural Networks. 2015 IEEE International Conference on Computer Vision (ICCV) 2015. p. 2956–64.

4. Yang X, Yan J, Wang W, Li S, Hu B, Lin J. Brain-inspired models for visual object recognition: an overview. Artificial Intelligence Review. 2022;55(7):5263–311.

5. Kanwisher N, Gupta P, Dobs K. CNNs reveal the computational implausibility of the expertise hypothesis. iScience. 2023;26(2).

6. Cadena SA, Denfield GH, Walker EY, Gatys LA, Tolias AS, Bethge M, et al. Deep convolutional models improve predictions of macaque V1 responses to natural images. PLOS Computational Biology. 2019;15(4):e1006897.

7. Bonner MF, Epstein RA. Computational mechanisms underlying cortical responses to the affordance properties of visual scenes. PLOS Computational Biology. 2018;14(4):e1006111.

8. Yamins DLK, Hong H, Cadieu CF, Solomon EA, Seibert D, DiCarlo JJ. Performance-optimized hierarchical models predict neural responses in higher visual cortex. Proceedings of the National Academy of Sciences of the United States of America 2014. p. 8619–24.

9. Mohsenzadeh Y, Mullin C, Lahner B, Oliva A. Emergence of Visual Center-Periphery Spatial Organization in Deep Convolutional Neural Networks. Scientific Reports. 2020;10(1).

10. Wallis TSA, Funke CM, Ecker AS, Gatys LA, Wichmann FA, Bethge M. A parametric texture model based on deep convolutional features closely matches texture appearance for humans. Journal of Vision. 2017;17(12):5.

11. Kuperwajs I, Schütt HH, Ma WJ. Using deep neural networks as a guide for modeling human planning. Scientific Reports. 2023;13(1).

12. Peterson JC, Abbott JT, Griffiths TL. Evaluating (and Improving) the Correspondence Between Deep Neural Networks and Human Representations. Cognitive Science. 2018;42(8):2648–69.

13. Jang H, McCormack D, Tong F. Noise-trained deep neural networks effectively predict human vision and its neural responses to challenging images. PLoS Biol. 2021;19(12):e3001418.

14. Kell AJE, Yamins DLK, Shook EN, Norman-Haignere SV, McDermott JH. A Task-Optimized Neural Network Replicates Human Auditory Behavior, Predicts Brain Responses, and Reveals a Cortical Processing Hierarchy. Neuron: Elsevier Inc.; 2018. p. 630–44.e16.

15. Ivet Rafegasa MV, Luís A. Alexandreb, Guillem Ariasa. Understanding Trained CNNs by Indexing Neuron Selectivity. 2019.

16. Ratan Murty NA, Bashivan P, Abate A, DiCarlo JJ, Kanwisher N. Computational models of category-selective brain regions enable high-throughput tests of selectivity. Nat Commun. 2021;12(1):5540.

17. VanRullen R. Reconstructing faces from fMRI patterns using deep generative neural networks. 2019.

18. Mcgugin RW, Van Gulick AE, Tamber-Rosenau BJ, Ross DA, Gauthier I. Expertise Effects in Face-Selective Areas are Robust to Clutter and Diverted Attention, but not to Competition. Cerebral Cortex. 2015;25(9):2610–22.

19. Bukach CM, Phillips WS, Gauthier I. Limits of generalization between categories and implications for theories of category specificity. Attention, Perception & Psychophysics. 2010;72(7):1865–74.

20. Brefczynski-Lewis JA, Lutz A, Schaefer HS, Levinson DB, Davidson RJ. Neural correlates of attentional expertise in long-term meditation practitioners. Proceedings of the National Academy of Sciences. 2007;104(27):11483–8.

21. Wong YK, Folstein JR, Gauthier I. The nature of experience determines object representations in the visual system. Journal of Experimental Psychology: General. 2012;141(4):682–98.

22. Zhang T, Dong M, Wang H, Jia R, Li F, Ni X, et al. Visual expertise modulates baseline brain activity: a preliminary resting-state fMRI study using expertise model of radiologists. BMC Neuroscience. 2022;23(1).

23. Gauthier I, Tarr MJ, Anderson AW, Skudlarski P, Gore JC. Activation of the middle fusiform ’face area’ increases with expertise in recognizing novel objects. Nature Neuroscience. 1999;2(6):568–73.

24. Wong AC-N, Palmeri TJ, Gauthier I. Conditions for Facelike Expertise With Objects. Psychological Science. 2009;20(9):1108–17.

25. Xu Y. Revisiting the Role of the Fusiform Face Area in Visual Expertise. Cerebral Cortex. 2005;15(8):1234–42.

26. Mcgugin RW, Newton AT, Gore JC, Gauthier I. Robust expertise effects in right FFA. Neuropsychologia. 2014;63:135–44.

27. Gauthier I, Skudlarski P, Gore JC, Anderson AW. Expertise for cars and birds recruits brain areas involved in face recognition. Nature Neuroscience. 2000;3(2):191–7.

28. Stokes D. On perceptual expertise. Mind & Language. 2021;36(2):241–63.

29. Harel A, Gilaie-Dotan S, Malach R, Bentin S. Top-Down Engagement Modulates the Neural Expressions of Visual Expertise. Cerebral Cortex. 2010;20(10):2304–18.

30. Kok EM, Sorger B, Van Geel K, Gegenfurtner A, Van Merriënboer JJG, Robben SGF, et al. Holistic processing only? The role of the right fusiform face area in radiological expertise. PLOS ONE. 2021;16(9):e0256849.

31. Martens F, Bulthé J, van Vliet C, Op de Beeck H. Domain-general and domain-specific neural changes underlying visual expertise. NeuroImage. 2018;169:80–93.

32. Bilalic M, Langner R, Ulrich R, Grodd W. Many Faces of Expertise: Fusiform Face Area in Chess Experts and Novices. Journal of Neuroscience. 2011;31(28):10206–14.

33. Stokes D. On perceptual expertise. Mind & Language. 2020;36(2):241–63.

34. Richler JJ, Wong YK, Gauthier I. Perceptual Expertise as a Shift From Strategic Interference to Automatic Holistic Processing. Current Directions in Psychological Science. 2011;20(2):129–34.

35. Kanwisher N, Gupta P, Dobs K. CNNs Reveal the Computational Implausibility of the Expertise Hypothesis. iScience. 2023.

36. Simonyan K, Zisserman A. Very deep convolutional networks for large-scale image recognition. arXiv preprint arXiv:14091556. 2014.

37. Deng J, Dong W, Socher R, Li L-J, Kai L, Li F-F. ImageNet: A large-scale hierarchical image database. 2009 IEEE Conference on Computer Vision and Pattern Recognition 2009. p. 248–55.

38. Parkhi OM, Vedaldi A, Zisserman A. Deep Face Recognition. Procedings of the British Machine Vision Conference 2015 2015. p. 41.1–.12.

39. Burge J, Geisler WS. Optimal defocus estimation in individual natural images. Proceedings of the National Academy of Sciences. 2011;108(40):16849–54.

40. Wennekers T, Dhamecha TI, Singh R, Vatsa M, Kumar A. Recognizing Disguised Faces: Human and Machine Evaluation. PLoS ONE. 2014;9(7).

41. Paterson K, Brodeur MB, Guérard K, Bouras M. Bank of Standardized Stimuli (BOSS) Phase II: 930 New Normative Photos. PLoS ONE. 2014;9(9).

42. Treue S, Trujillo JCM. Feature-based attention influences motion processing gain in macaque visual cortex. Nature. 1999;399(6736):575–9.

43. McAdams CJ, Maunsell JHR. Effects of Attention on Orientation-Tuning Functions of Single Neurons in Macaque Cortical Area V4. The Journal of Neuroscience. 1999;19(1):431–41.

44. Lee J, Maunsell JHR. The Effect of Attention on Neuronal Responses to High and Low Contrast Stimuli. Journal of Neurophysiology. 2010;104(2):960–71.

45. Compte A, Wang X-J. Tuning Curve Shift by Attention Modulation in Cortical Neurons: a Computational Study of its Mechanisms. Cerebral Cortex. 2006;16(6):761–78.

46. Haenny PE, Schiller PH. State dependent activity in monkey visual cortex. Experimental Brain Research. 1988;69(2):225–44.

47. Doostani N, Hossein-Zadeh G-A, Cichy RM, Vaziri-Pashkam M. Attention Modulates Human Visual Responses to Objects by Tuning Sharpening. 2023.

48. Ling S, Liu T, Carrasco M. How spatial and feature-based attention affect the gain and tuning of population responses. Vision Research. 2009;49(10):1194–204.

49. Cohen MA, Konkle T, Rhee JY, Nakayama K, Alvarez GA. Processing multiple visual objects is limited by overlap in neural channels. Proceedings of the National Academy of Sciences. 2014;111(24):8955–60.

50. Kiat JE, Luck SJ, Beckner AG, Hayes TR, Pomaranski KI, Henderson JM, et al. Linking patterns of infant eye movements to a neural network model of the ventral stream using representational similarity analysis. Developmental Science. 2022;25(1).

51. Diedrichsen J, Khaligh-Razavi S-M, Kriegeskorte N. Deep Supervised, but Not Unsupervised, Models May Explain IT Cortical Representation. PLoS Computational Biology. 2014;10(11).

52. Peelen MV, Kastner S. A neural basis for real-world visual search in human occipitotemporal cortex. Proceedings of the National Academy of Sciences. 2011;108(29):12125–30.

53. Noah S, Powell T, Khodayari N, Olivan D, Ding M, Mangun GR. Neural Mechanisms of Attentional Control for Objects: Decoding EEG Alpha When Anticipating Faces, Scenes, and Tools. Journal of Neuroscience 2020. p. 4913–24.

54. Folstein JR, Monfared SS, Maravel T. The effect of category learning on visual attention and visual representation. Psychophysiology. 2017;54(12):1855–71.

55. Reynolds JH, Chelazzi L. Attentional Modulation of Visual Processing. Annual Review of Neuroscience. 2004;27(1):611–47.

56. Grabner RH, Neubauer AC, Stern E. Superior performance and neural efficiency: The impact of intelligence and expertise. Brain Research Bulletin. 2006;69(4):422–39.

57. Maguire EA, Valentine ER, Wilding JM, Kapur N. Routes to remembering: the brains behind superior memory. Nature Neuroscience. 2002;6(1):90–5.

58. Wolfe JM. Guided Search 2.0 A revised model of visual search. Psychonomic Bulletin & Review. 1994;1(2):202–38.

59. Williams M, Becker SI. Determinants of Dwell Time in Visual Search: Similarity or Perceptual Difficulty? PLoS ONE. 2011;6(3).

60. Hout MC, Goldinger SD. Target templates: the precision of mental representations affects attentional guidance and decision-making in visual search. Attention, Perception, & Psychophysics. 2014;77(1):128–49.

61. Lee J, Geng JJ. Idiosyncratic Patterns of Representational Similarity in Prefrontal Cortex Predict Attentional Performance. The Journal of Neuroscience. 2017;37(5):1257–68.

62. Sabine Kastner MAP, Peter De Weerd, Robert Desimone, and Leslie G. Ungerleider. Increased Activity in Human Visual Cortex during Directed Attention in the Absence of Visual Stimulation. 1999.

63. Reynolds JH, Chelazzi L, Desimone R. Competitive Mechanisms Subserve Attention in Macaque Areas V2 and V4. The Journal of Neuroscience. 1999;19(5):1736–53.

64. Bukach CM, Gauthier I, Tarr MJ. Beyond faces and modularity: the power of an expertise framework. Trends in Cognitive Sciences. 2006;10(4):159–66.

65. van der Linden M, Wegman J, Fernández G. Task- and Experience-dependent Cortical Selectivity to Features Informative for Categorization. Journal of Cognitive Neuroscience. 2014;26(2):319–33.

66. Reddy L, Kanwisher N. Category Selectivity in the Ventral Visual Pathway Confers Robustness to Clutter and Diverted Attention. Current Biology. 2007;17(23):2067–72.

67. Zhang W, Luck SJ. Feature-based attention modulates feedforward visual processing. Nature Neuroscience. 2008;12(1):24–5.

